# Lipid Landscape of human cells

**DOI:** 10.1101/2025.10.05.680593

**Authors:** Mads Møller Foged, Knut Kristoffer Bundgaard Clemmensen, Lya Katrine Kauffeldt Holland, Jano Dicroce Giacobini, Mesut Bilgin, Kevin Titeca, Anne-Claude Gavin, Marja Jäättelä, Kenji Maeda

## Abstract

Eukaryotic cells synthesise thousands of lipid species, yet the principles that organise them across compartments remain unclear. Here we introduce PAP-SL, parallel (immuno)affinity purification coupled to shotgun lipidomics, to map ∼300 lipid species across six compartments of human cells and uncover general rules of lipid organisation. Correlation analyses of stoichiometric lipid compositions resolved archetypal distribution trajectories. Biosynthetic origin set the baseline: mitochondrial and lysosomal pathways generated confined pools, whereas the ER acted as a dispatch hub shaping downstream membrane territories. The ER established the saturated plasma membrane through transfer of phosphatidylserine, sphingolipids, and cholesterol, together with species-level selection of more saturated glycerophospholipids across diverse headgroups, while less-saturated species moved unselectively into mitochondria. The ER also channelled ether-linked glycerophospholipids to lysosomes, forming a territory dominated by ether phosphatidylcholine that collapsed under ethanolamine-driven metabolic rewiring. PAP-SL defines a framework to resolve lipid organisation and its metabolic plasticity in health and disease.

## Introduction

Lipids are extraordinary in their diversity. Human cells synthesize thousands of distinct molecular species, differing in backbone, headgroup, hydrocarbon chain type, length, unsaturation, and linkage to the backbone^1–4^. This structural diversity underlies a striking compositional heterogeneity across compartments, each tailored to their architecture and functions^3,5,6^. Yet we still lack a cell-wide view of how individual lipid species are distributed or what principles organise them^3^.

This compositional heterogeneity is fundamental to membrane function. It creates a varied membrane landscape in which local mechanics and protein interactions are finely tuned: matching transmembrane helices to bilayer thickness, recruiting peripheral proteins through electrostatic and structural cues. These together provide the spatiotemporal control needed for orchestration of cellular processes^7–10^. Mitochondria rely on cardiolipin (CL) to maintain cristae architecture and respiratory efficiency^11^, whereas the late endosomes and lysosomes (together referred to as lysosomes from here on) depend on bis(monoacylglycero)phosphate (BMP) to drive catabolism^12,13^. Beyond these examples of compartmentalized biosynthesis, cholesterol, sphingomyelin (SM), and phosphatidylserine (PS) form steep concentration gradients along the secretory pathway, establishing a thick and robust plasma membrane (PM) that supports protein network assembly for signal transduction^6,14^. These gradients reflect the underlying organisation of lipid metabolism and transport, centred on the endoplasmic reticulum (ER) as the main production hub. Local conversions and both vesicular and non-vesicular trafficking of ER-derived lipids diversify lipid compositions across compartments, aided by lipid transfer proteins (LTPs) that mediate direct exchange between membranes. Among these, oxysterol-binding protein–related proteins (ORPs) shuttle ER-synthesised cholesterol and PS to distal compartments^14–17^. Thus, cellular membrane landscapes emerge from coordinated lipid synthesis, conversion, and transport, and their breakdown contributes to human disease^18,19^.

Despite these paradigmatic examples, current knowledge of lipid organization remains fragmentary. Most insights derive from a few lipid classes studied in selected compartments, whereas the broader distribution of individual species, which share biosynthetic origins but differing in hydrocarbon chain composition, has scarcely been explored. A systematic, cell-wide view of how the lipidome partitions across compartments is still lacking, largely because existing approaches fall short: shotgun lipidomics provides a stoichiometric and broad coverage of the lipidome, but usually at whole-cell level or, in specialized cases, from isolated organelles one at a time^17,20–22^. In contrast, probe-based imaging tracks only a limited set of lipid classes such as PS and cholesterol^23–25^. Without a comprehensive framework, the principles that govern lipid distribution remain unresolved.

To overcome this gap, we developed PAP-SL—parallel (immuno)affinity purification coupled with quantitative shotgun lipidomics—enabling simultaneous isolation and analysis of multiple compartments from the same cells. Applied to human cells, PAP-SL generated stoichiometric lipidomes for six compartments, resolving the distributions of hundreds of lipid species. These maps uncover general principles of lipid organisation and establish a framework for exploring how lipid landscapes are regulated in physiology and disease.

## Results

### PAP-SL provides quantitative, compartment-resolved maps of the cellular lipidome

We established PAP-SL in HeLa cells. PM was affinity purified by cell-surface biotinylation of intact cells followed by capture from a post-nuclear fraction (PNF) on streptavidin-conjugated paramagnetic microbeads^21^ (Fig. 1a). In parallel, the same PNF preparation was divided into matched aliquots, which were subjected to immunoaffinity purification of additional compartments using antibodies against endogenous markers: TOM20 (mitochondria), calnexin (ER), RCAS1 (cis-Golgi), TGN46 (trans-Golgi network, TGN), LAMP1 (lysosomes)—or control IgG, and secondary antibody–conjugated paramagnetic microbeads^17,20–22^. Peroxisomes (PEX14) were tested but excluded from the final protocol (see below). Specificity was validated by Western blotting with only minor cross-contamination (Extended Data Fig. 1).

**Figure 1.**
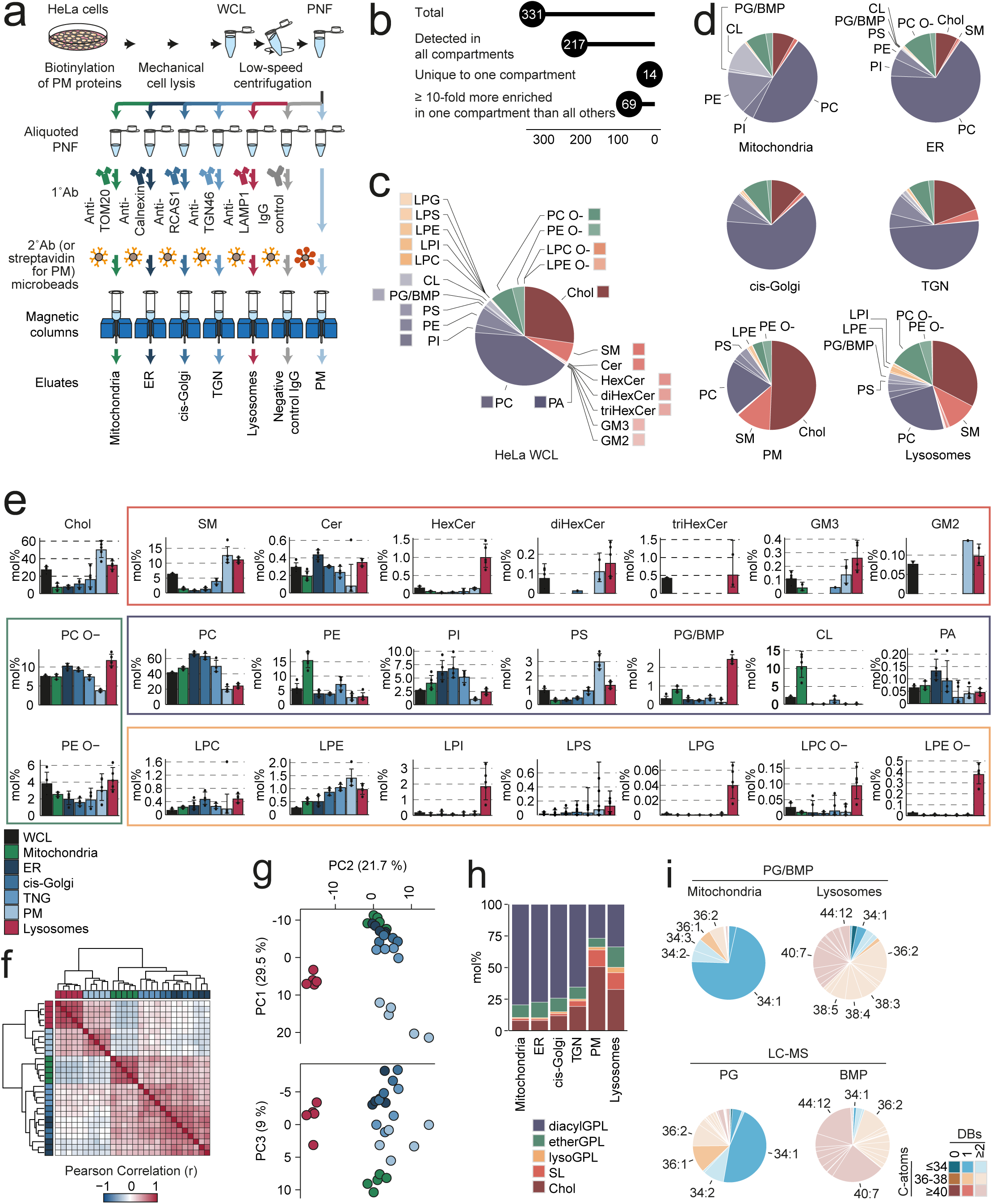
PAP-SL enables stoichiometric mapping of ∼300 lipid species across six compartments. **a**, Schematic of the PAP-SL workflow. **b**, Total lipid species in the dataset and their distribution across compartments. **c**, mol% of lipid classes in whole-cell lysate (WCL). **d–e**, mol% of lipid classes in isolated compartments, shown as pie charts by compartment (**d**) or bar graphs by class (**e**). **f**, Hierarchical clustering of compartments using Pearson correlation coefficients of log_2_ mol% values across all species. **g**, Principal component analysis of compartments based on log_2_ mol% values. **h**, mol% of indicated lipid (sub)categories across compartments. **i**, Species composition of PG/BMP in mitochondria and lysosomes determined by shotgun lipidomics (top) and of PG and BMP in WCL determined by LC–MS (bottom). Abbreviations: ER, endoplasmic reticulum; IgG Ctrl, isotype control; Ab, antibody; BMP, bis(monoacylglycero)phosphate; Cer, ceramide; Chol, cholesterol; CL, cardiolipin; diHexCer, dihexosylceramide; etherGPL, ether glycerophospholipid; GM2, ganglioside GM2; GM3, ganglioside GM3; GPL, glycerophospholipid; HexCer, hexosylceramide; LPC, lysoPC; LPC O–, alkyl LPC; LPE, lysoPE; LPE O–, alkyl LPE; LPG, lysoPG; LPI, lysoPI; LPS, lysoPS; lysoGPL, lysoglycerophospholipid; PA, phosphatidic acid; PC, phosphatidylcholine; PC O–, alkyl-acyl PC; PE, phosphatidylethanolamine; PE O–, alkyl-acyl PE; PG, phosphatidylglycerol; PI, phosphatidylinositol; PS, phosphatidylserine; SL, sphingolipid; SM, sphingomyelin; triHexCer, trihexosylceramide.

We then absolutely quantified lipids in fractions from five independent PAP rounds using shotgun lipidomics on a high-resolution MS^26^, with spike-in standards (Extended Data Table 1). Ions were assigned to lipid species defined by class and sum composition (e.g., PS 34:1, see Methods) using class-specific identification criteria (Extended Data Table 2) and filtered against blanks and databases (see Methods, overview in Extended Data Fig. 2). We excluded seven lipid classes with unstable quantification across replicates, and peroxisomes due to low yields indistinguishable from IgG controls (Extended Data Fig. 3a). The final HeLa PAP-SL dataset comprised 331 lipid species across 24 classes: cholesterol, seven sphingolipid (SL) classes, and sixteen glycerophospholipid (GPL) classes. Among GPLs, acyl-acyl GPL (diacylGPL; including CL and BMP, which carry atypical acyl configurations), alkyl-acyl GPL (etherGPL), and monoacyl/alkyl GPL (lysoGPL) were treated as separate classes.

Most of these lipid classes were detected across multiple compartments (Fig. 1b). We normalized abundances to molar percentage (mol%) of total lipids to enable direct comparisons of lipid compositions despite variation in total lipid yields between compartments and replicates (Extended Data Fig. 3b)^27^. This revealed HeLa whole-cell lysate (WCL) dominated by cholesterol (27 mol%), phosphatidylcholine (PC) (41 mol%), and SM (6.3 mol%) (Fig. 1c). In comparison, each isolated compartment displayed a unique lipid composition (Fig. 1d).

Correlation and principal component analyses of these lipid compositions grouped the compartments into two sets: ER, cis-Golgi, TGN, and mitochondria (slightly offset) versus PM and lysosomes (Fig. 1f,g). A steep cholesterol gradient from 7.8 mol% in the ER to 50 mol% at the PM accompanied this division (Fig. 1h), consistent with ORP-mediated export of cholesterol after ER synthesis^15^. In parallel, total diacylGPLs fell from 77 mol% in the ER to 26 mol% at the PM. Together, these features reflect the established concept that cellular membranes are organized into two territories separated by the secretory pathway^7^.

Within these two territories, mitochondria and lysosomes stood out with distinctive signatures reflecting local lipid biosynthesis: CL, phosphatidylethanolamine (PE), and phosphatidylglycerol (PG) in mitochondria, and BMP in multivesicular bodies^5,11,28–30^. CL was enriched from 1.9 mol% in the WCL to 10.5 mol% in mitochondria but was virtually absent elsewhere, with only a minor signal in the TGN (1.2 mol%) (Fig. 1e). PE was likewise enriched from 5.6 mol% in the WCL to 15.4 mol% in mitochondria, where it is produced by decarboxylation of PS headgroup. In contrast, PE reached only 3.8 mol% in the ER despite parallel synthesis through the Kennedy pathway, which transfers phosphoethanolamine headgroup from cytidine diphosphate (CDP)–ethanolamine onto diacylglycerol (DAG) (Fig. 1e).

Because our shotgun lipidomics setup cannot distinguish BMP from its structural isomer PG, PAP-SL reported them jointly here as PG/BMP (quantified using a PG standard)^26^. Even so, PG/BMP was enriched from just 0.32 mol% in the WCL to 2.4 mol% in lysosomes, while it was depleted from the ER (0.26 mol%) and PM (0.12 mol%) (Fig. 1e). Importantly, this lysosomal PG/BMP pool almost entirely consisted of species having ≥2 double bonds (DBs); a feature that matched BMP in HeLa WCL and contrasted with predominantly monounsaturated PG, as resolved by LC–MS (Fig. 1i). Mitochondria were also enriched in PG/BMP to 0.82 mol% (Fig. 1e). This pool was dominated by PG/BMP 34:1 (73 mol% of the class), consistent with local PG biosynthesis (Fig. 1i).

Altogether, these data show that PAP-SL accurately resolves compartment-specific lipid compositions, capturing both established gradients and locally synthesized pools, with minor cross-contamination confined to mitochondrial lipids in the TGN.

### Lipid co-localization networks uncover six archetypal modes of organization

With compartmental lipid compositions validated, we next moved on to generate an atlas of lipids and elucidate rules governing their partitioning. Relative to WCL, most lipid species showed relative enrichment in some compartments and depletion in others, consistent with genuine spatial partitioning rather than systematic bias (Fig. 2a). Species within the same class often diverged, highlighting the need to analyze distributions at the species level rather than relying on aggregated class profiles.

**Figure 2.**
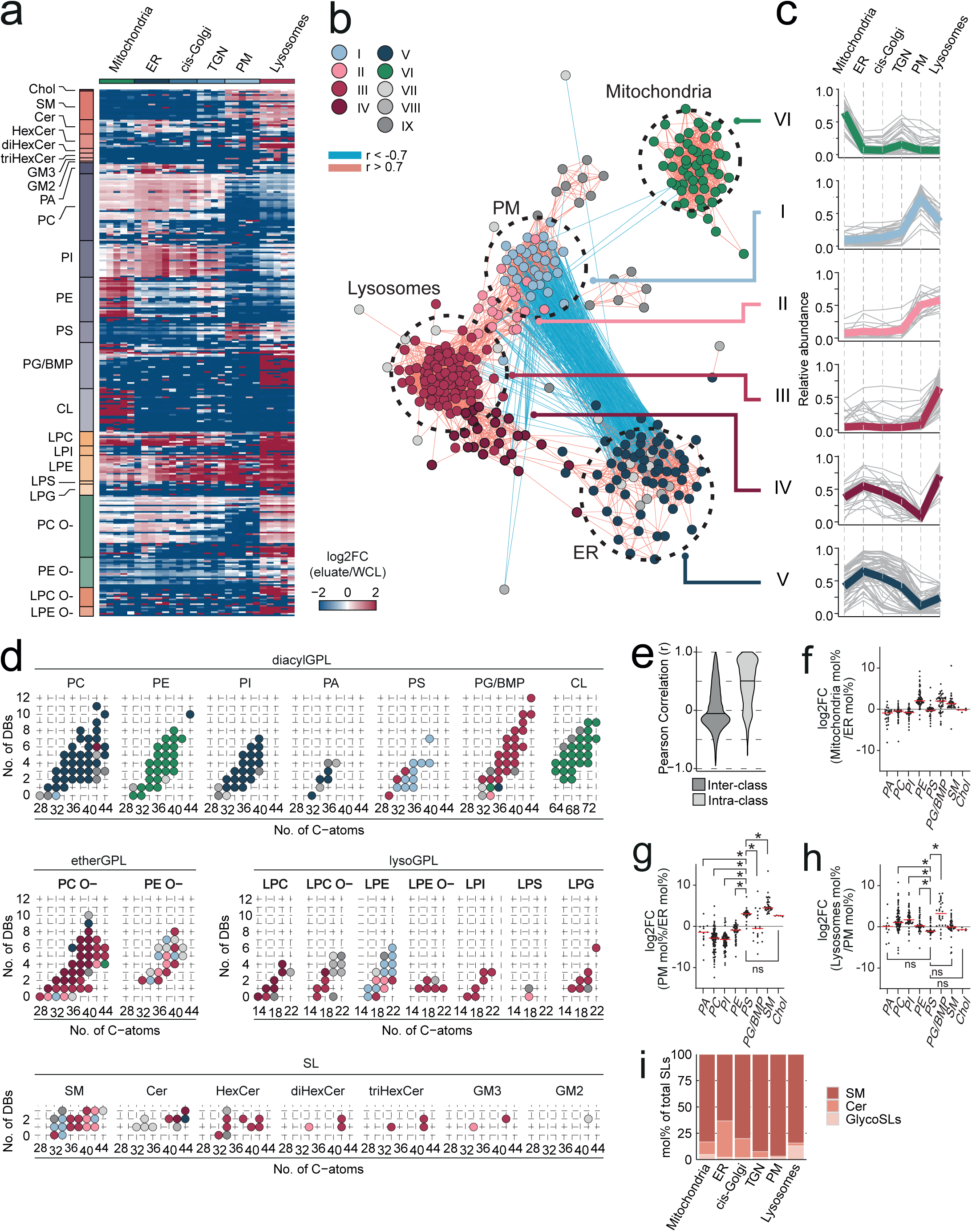
PAP-SL reveals lipid co-localization networks. **a**, Log_2_-transformed fold changes (FC) in mol% of lipid species across compartments relative to whole-cell lysate (WCL). **b**, Lipid–lipid correlation network (Pearson correlation coefficient (r); red: >0.7, blue: <–0.7) based on species mol% (of total excluding cholesterol). Species are coloured by hierarchical clustering. Non-correlated species (|r|<0.7) were excluded. **c**, Cluster mean distribution profiles of lipid species across compartments (grey: individual species; coloured: cluster means, normalized to maximum mol%). **d**, Species-to-cluster assignments grouped by class and (sub)category, arranged by DBs (vertical) and total carbons (C). **e**, Violin plots of correlation distributions for lipid pairs within or between classes. **f–h**, Species-level log2 fold changes for ER to mitochondria (**f**), ER to PM (**g**), and PM to lysosomes (**h**) (dots = replicates, red = medians). Significance from Welch’s ANOVA with Benjamini–Hochberg correction (FDR 1%). **i**, mol% of SM, total glycoSLs, and Cer within the total SL pool across compartments. *N = 4–5. Abbreviations as in* Fig. 1.

To systematically capture recurring distribution patterns among species, we computed pairwise Pearson correlations using their mol% values across compartments (excluding cholesterol in mol% calculation to avoid overweighting). Hierarchical clustering revealed nine clusters of co-distributed species (Extended Data Fig. 4). In parallel, we constructed a correlation network restricted to the strong positive or negative correlations (|r|>0.7). Both approaches converged on six clusters (Clusters I–VI; 261 lipid species), which emerged as discrete modules in the network (Fig. 2b).

These six modules defined archetypal modes of lipid distribution, as captured by their cluster-mean profiles (Fig. 2c). Four modules showed a clear peak in a single compartment: ER (Cluster V; ER module), mitochondria (Cluster VI; mitochondrial module), PM (Cluster I; PM module), and lysosomes (Cluster III; lysosomal module), and thus served as compartmental landmarks in the network (Fig. 2b). The remaining two modules occupied intermediate positions, with dual maxima bridging PM with lysosome (Cluster II) and ER with lysosomes (Cluster IV), respectively (Fig. 2b,c).

Species-to-cluster assignments revealed strong intra-class coherence within most classes (Fig. 2d) and overall (median r = 0.52 vs –0.057 for inter-class) (Fig. 2e), showing that species sharing a common biosynthetic origin tended to co-distribute.

DiacylGPL classes mapped to the modules of their biosynthetic sites (Fig. 2d):PC, phosphatidylinositol (PI), and phosphatidic acid (PA) to the ER module (Cluster V), CL, PE, and PG/BMP 34:1 (representing PG) to the mitochondrial module (Cluster VI), and PG/BMP species with ≥2 DBs (representing BMP) to the lysosomal module (Cluster III)^6^—indicating that biosynthetic origin determines their primary localization.

### LTP-mediated transfer overrides origin-based lipid trajectories

Lipids in mitochondrial and lysosomal modules appeared largely confined to their biosynthetic sites (Fig. 1e, 2c). By contrast, lipids of ER origin dispersed more broadly.

PC, PI, and PA peaked in the ER and declined only modestly in mitochondria (Fig. 2f) (PC 47.5 mol% vs 66.2 mol% at ER; Fig. 1e) consistent with bulk flow. Their abundances dropped more sharply along the secretory pathway (Fig. 2g), yet they persisted at considerable levels at the PM (PC 19.6 mol%) and even rebounded at lysosomes (Fig. 2h) (PC 24.2 mol%). These parallel trajectories of PC, PI, PA, and even PE (Fig. 2g,h) indicate that GPLs of various headgroups can follow similar trajectories.

PS, an ORP-cargo^14,31,32^, broke this pattern. Despite its ER origin, it mapped to the PM module (Cluster I). Its abundance rose sharply from 0.31 mol% at the ER to 3.0 mol% at the PM (Fig. 1e, 2g) and then declined toward lysosomes (1.3 mol%) (Fig. 2h). The PM-focused distribution of PS mirrored that of another ORP cargo, cholesterol^15^ (Fig. 2g,h), underscoring that LTP-mediated transport is central to establishing the lipid identity of the PM.

SLs reinforced this conclusion. The dominant SM and more minor glycosphingolipid (glycoSL; HexCer, diHexCer, triHexCer, GM3, GM2) classes primarily clustered to modules associated with PM and lysosomes (Fig. 2d; Clusters I, II, and III). In contrast, their precursor, ceramide, showed weak cluster assignment and a rather flat distribution across compartments (Fig. 1e). Despite being synthesized in the ER and transferred to Golgi by CERT, ceramide itself did not accumulate along the secretory pathway, unlike PS and cholesterol. Instead, its transfer coincided with expansion of the total SL pool from 1.3 mol% in the ER to 13 mol% at the PM (Fig. 1h)—driven by a rise in ceramide-derived SM (0.79 mol% in the ER to 12.6 mol% at the PM; Fig. 1e). Consistent with conversion at the Golgi, ceramide’s share of the total SL pool dropped from 34.4 mol% in the ER to only 0.3 mol% at the PM. GlycoSLs followed distributions similar to SM but with stronger lysosomal polarization. Together, these observations highlight a mechanism in which transfer of an ER-synthesized precursor is coupled to downstream synthesis, expanding the SL pool at the PM.

These examples show that ER-synthesized lipids either follow origin-based trajectories, remaining abundant in the ER and also moving in bulk to mitochondria, or escape this fate through selective transfer by LTPs such as ORPs and CERT to establish PM composition.

### EtherGPLs break origin rules and establish a lysosomal territory of non-canonical GPLs

Above, we showed that headgroup-specific transfer (e.g., PS via ORPs) can divert ER-synthesized GPLs from broad, origin-based trajectories to a PM-focused distribution. Here, we find that chemical linkage can do the same: an sn-1 ether bond redirects ER-made GPLs toward a lysosomal territory where conventional diacylGPLs are relatively scarce.

Ether PC (PC O–) and ether PE (PE O–) carry an sn-1 ether-linked fatty alcohol (Falc) yet share the Kennedy pathway with PC and PE, using alkyl-DAG rather than DAG as the headgroup acceptor (Fig. 3a)^33^. Despite their ER origin, the two ether classes diverged strongly in their compartmental trajectories from their diacyl counterparts and from one another.

**Figure 3.**
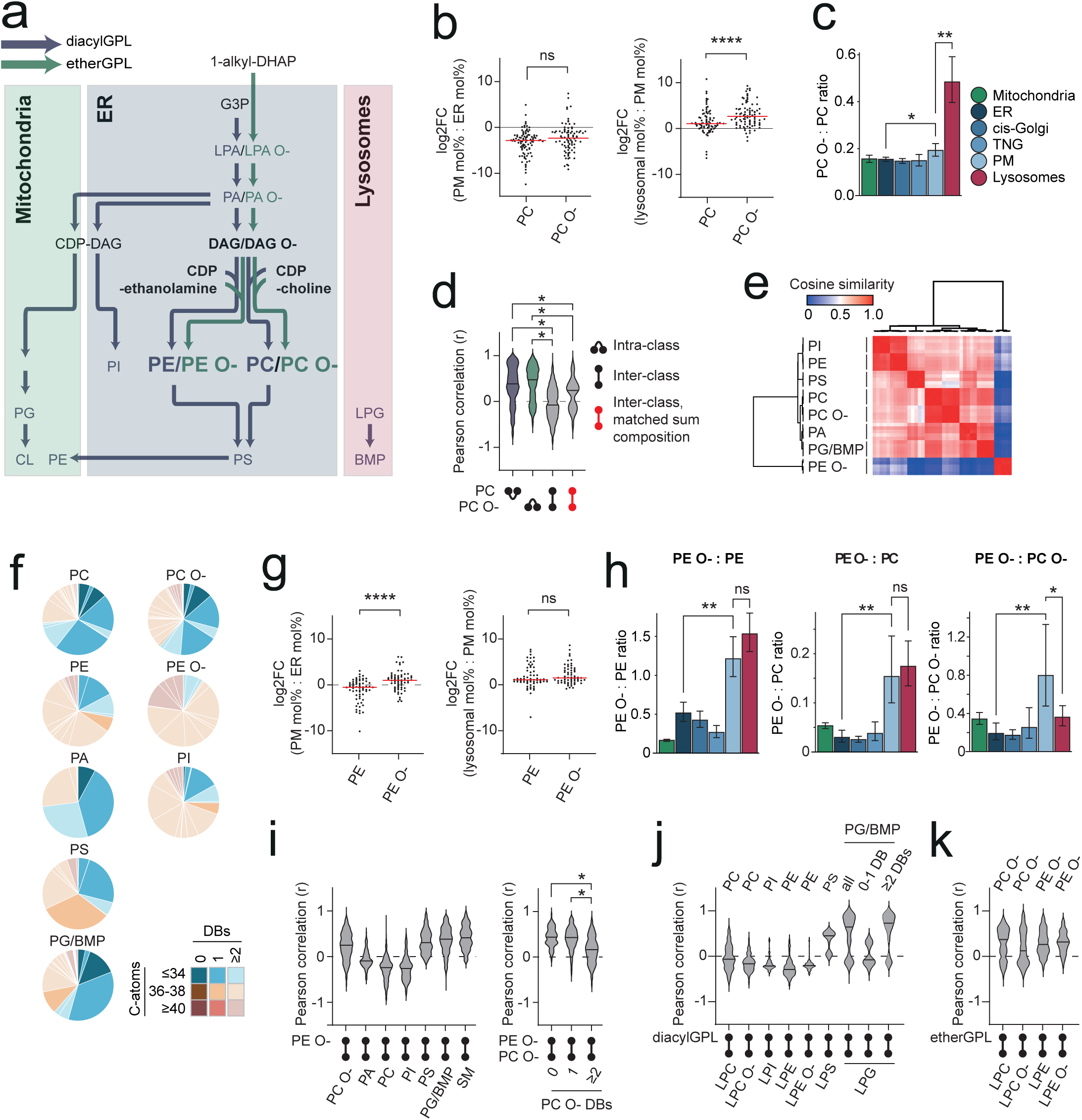
PAP-SL uncovers a lysosomal territory enriched in ether GPLs. **a**, Schematic of de novo glycerophospholipid biosynthetic pathways mapped to compartments. **b**, Log_2_-transformed fold changes (FC) of PC and PC O– species for ER to PM (left) and PM to lysosomes (right) (dots = replicates, red = median). **c**, PC O–:PC class ratios across compartments. **d**, Pearson correlation (r) for species within or between PC and PC O– classes, with or without matched sum compositions. **e**, Hierarchical clustering of GPL classes based on cosine similarity matrix of species compositions (mol% within class, whole-cell lysate). **f**, Species composition of indicated lipid classes, coloured by hydrocarbon chain double bond (DB) and carbon (C) counts. **g**, Log_2_-transformed FC of PE and PE O– species for ER to PM (left) and PM to lysosomes (right) (dots = replicates, red = median). **h**, PE O–:PE (left), PE O–:PC (middle), and PE O–:PC O– (right) class ratios across compartments. Colours are as in (**c**). **i**, Pearson correlations (r) between species of PE O– vs other classes (left) or PC O– species stratified by DB counts (right). **j-k**, Pearson correlations (r) between species of lysoGPLs and diacylGPLs (**j**) or lysoGPLs and etherGPLs (**k**). Statistics: Welch’s ANOVA with BH correction (FDR 1%) for **d** and **i**; Welch’s t-test for **b, c, g,** and **h**. *P*<0.05, \**P*<0.01, \*\**P*<0.001, \*\*\**P*<0.0001. *N = 4–5. Abbreviations as in* Fig. 1.

PC O– mapped to the ER–lysosome module (Cluster IV; Fig. 2d) and displayed dual peaks in lysosomes and the ER (Fig. 1e). Its abundance declined from 10.2 mol% in the ER to 3.8 mol% at the PM (Fig. 3b), while maintaining a stable PC O–:PC ratio of ∼0.15 (Fig. 3c). Beyond the PM, however, PC O-rose more steeply than PC toward lysosomes (11.7 mol%) (Fig. 3b), while elevating the PC O–:PC ratio up to 0.5 (Fig. 3c), suggesting that ether linkage caused lysosomal accumulation of PC O-.

This divergence of PC O-from PC is best attributed to the ether linkage itself rather than possible variations in other hydrocarbon chain features: Pearson correlations of compartmental distribution remained weak even for matched sum compositions (e.g., PC O– 34:1 vs PC 34:1) (Fig. 3d), and in HeLa WCL the two classes had highly similar species compositions—closer to each other than to any other GPLs (Fig. 3e,f).

PE O– clustered with PM- and lysosome-associated modules (Clusters I–III) (Fig. 2d). Its abundance rose modestly from the ER (2.0 mol%) toward the PM (3.0 mol%) and lysosomes (4.2 mol%; Fig. 1e). This contrasted with the general decline of most ER-synthesized diacylGPLs along the secretory pathway (Figs. 2g and 3g), resulting in relative enrichment of PE O– compared with PE, PC, and even PC O– (Fig. 3h). From PM to lysosomes, however, PE O– tracked similarly to PE and PC, diverging only relative to PC O– (Fig. 3h). Thus, while both ether classes peaked in lysosomes, PC O– did so by accumulating specifically in lysosomes, whereas PE O– largely carried forward abundance gained at the PM.

PE O– consisted entirely of species with ≥2 DBs and thereby greatly differed from PC O- and diacyl GPL classes in hydrocarbon chain features (Fig. 3e,f). Yet unsaturation alone cannot explain its trajectory, since even PC O– species with ≥2 DBs correlated poorly in compartmental distribution with PE O– (Fig. 3i).

Together, these findings establish that ether linkage overrides origin-based trajectories and concentrate etherGPLs in lysosomes. However, the mechanisms underlying this targeting remain to be clarified.

Besides etherGPLs, lysoGPLs were likewise concentrated at lysosomes, consistent with their catabolic function. Most lysoGPL species clustered to the lysosomal module (Cluster III; Fig. 2d) and correlated poorly with their diacyl and ether counterparts, except for the BMP precursor LPG^28–30^ with PG/BMP (Fig. 3j,k). Class-level distribution profiles were consistent, showing lysosome-centered distributions (Fig. 1e), in line with lysosomal catabolic activity.

Alongside etherGPLs, lysoGPLs further reshaped the lysosomal GPL landscape: etherGPLs expanded to 30% of total GPLs (from ∼12–14% in ER/Golgi) and lysoGPLs to 8% (from ∼1–2%), underscoring lysosomes as a distinct membrane territory enriched in non-canonical GPLs, with PC O– as the dominant class.

### Dual strategies establish PM saturation

So far, PAP-SL revealed that ER-synthesized GPLs reach well beyond the ER, with distributions dictated by headgroup identity (e.g., PS vs others) and ether linkage (e.g., PC O- vs PC). However, many species deviated from these class-wise trajectories, revealing additional species-level principles that further govern lipid distributions.

Fully saturated PC and PC O– species were prominent examples. They tended to cluster with the PM module (Cluster I)(Fig. 2d) and segregated from more unsaturated counterparts in the network (Fig. 4a). Unlike unsaturated species, which followed the class-wise trend of declining from the ER to the PM and rebounding in lysosomes, saturated PC species remained relatively stable across compartments (Fig. 4e,f). This selective behaviour shifted species composition of PC class: fully saturated PC species accounted for 31 mol% of the class at the PM, compared to 9 mol% in the ER and 16 mol% in lysosomes (Fig. 4d).

**Figure 4.**
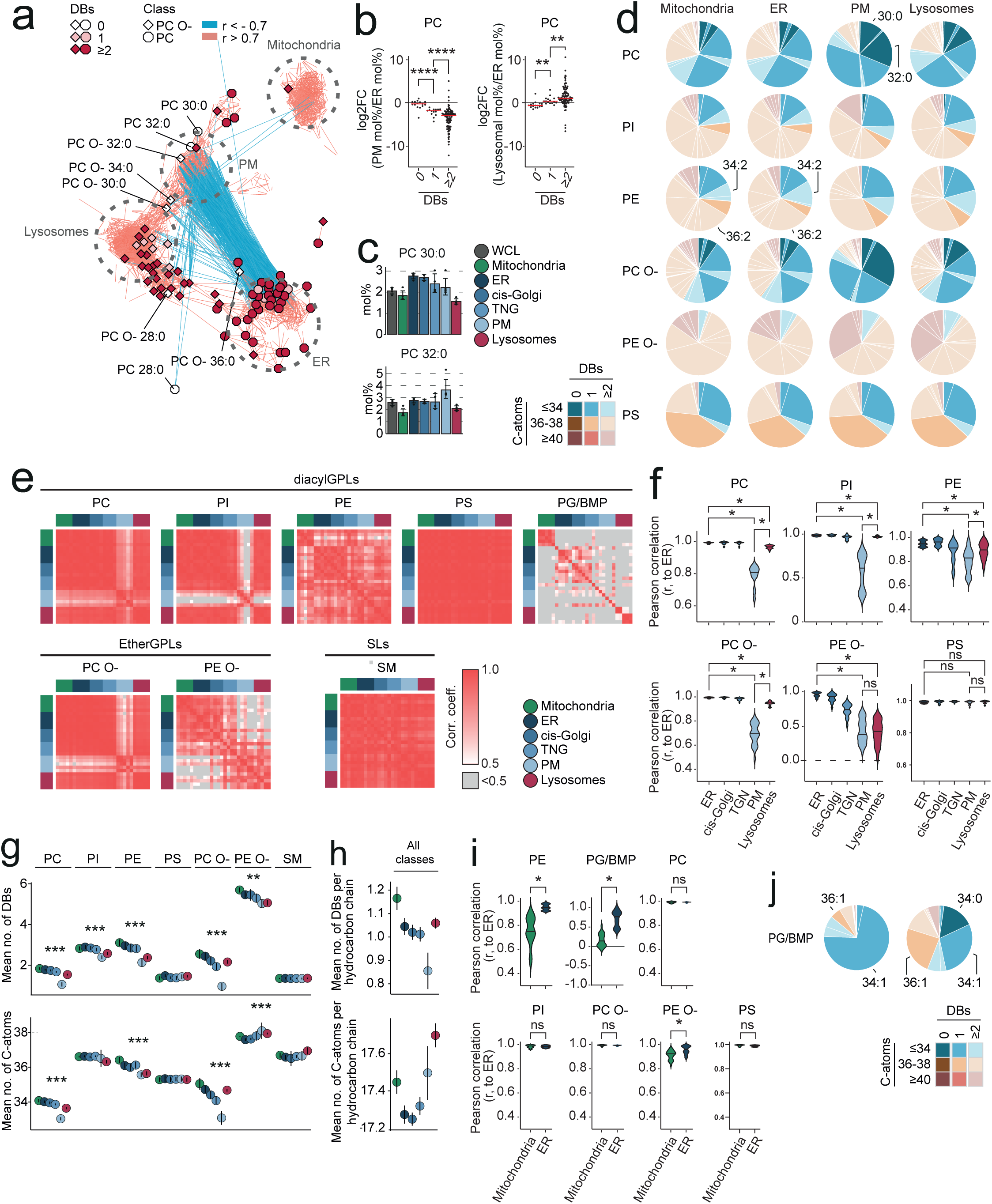
PAP-SL reveals PM-focused saturation and ER–mitochondria divergence. **a**, PC and PC O– species mapped onto the lipid network from Fig. 2b, coloured by hydrocarbon chain double (DB) and carbon (C) counts. **b**, Log_2_-transformed fold changes (FC) of PC species for ER to PM (left) and PM to lysosomes (right) (dots = replicates, red = median). **c**, mol% of PC 30:0 and PC 32:0 across compartments (mean ± SD). **d**, Species compositions (mol% within class) of indicated classes across compartments. **e**, Correlation matrices of species compositions across compartments for indicated classes. **f**, Pearson correlation coefficients (r) within ER or between ER and indicated compartments. **g**, Mean DBs (top) and C atoms (bottom) per species for major classes across compartments (mean ± SD). **h**, Mean DBs (top) and C atoms (bottom) per hydrocarbon chain across all lipid species (excluding cholesterol). **i**, Pearson correlation coefficients within ER or between ER and mitochondria. **j**, Species compositions (mol% within class) for PG/BMP in mitochondria vs ER. Statistics: Welch’s ANOVA with BH correction (FDR 1%) for **f** and **i**; Welch’s t-test for **b**; one-way ANOVAs carried out independently for each class with the null-hypothesis of no difference in means between compartments with Holm method correction for **g**. *P*<0.05, \**P*<0.01, \*\**P*<0.001, \*\*\**P*<0.0001. *N = 4–5. Abbreviations as in* Fig. 1.

While several other species deviated from class-wise trends in species-to-cluster assignment, many of them likely reflected poor measurements due to low abundance or noise. Moreover, the clustering approach may have missed genuine species-level deviations in distribution patterns that were not strong enough to alter cluster assignment. To identify deviations that systematically influence lipid organization, we therefore next followed the example of PC and searched for compartment-specific alterations in species compositions within each GPL class.

This analysis revealed two recurring organizational patterns among GPL classes (Fig. 4b). The first extended the principle seen for PC to essentially all other ER-synthesized GPL classes with exception of PS: they displayed a markedly different species composition at the PM compared with other compartments (Fig. 4b). In particular, the divergence from the ER (Fig. 4c) indicates that GPL classes are reshaped in species compositions as they progress along the secretory pathway. By contrast, lysosomal species compositions remained more similar to those of the ER, though to varying degrees (Fig. 4c). Notably, these trends were absent for the ORP cargo PS and for Golgi-synthesized SM, both of which maintained consistent compositions across compartments (Fig. 4b).

Although each GPL class displayed distinct species compositions at their origin and manners of reshaping in other compartments, they converged on a shared trend: increased saturation at the PM. PC exemplified this, showing a gradual drop in mean DBs per molecule from 1.8 in the ER to 1.1 at the PM, with partial reversal to 1.5 in lysosomes (Fig. 4g). This trend was evident across ER-synthesized GPLs, but not in PS or SM, which showed little change across compartments. Instead, these PM-localized classes were intrinsically more saturated than others, with mean DBs of ∼1.3–1.4 per molecule (Fig. 4g).

The PM also showed enhanced saturation when viewed at the level of entire lipidome: mean DBs per hydrocarbon chain. calculated for all quantified lipids excluding cholesterol, progressively declined from ER to PM and rebounded in lysosomes, while mean C-atoms steadily increased across these compartments (Fig. 4h).

Together, these findings point to two complementary selection principles underlying PM saturation: whole-class recruitment of intrinsically saturated PS and SM, and enrichment of more saturated species within otherwise unsaturated, poorly PM-represented classes.

### Asymmetric lipid trafficking between ER and mitochondria

The systematic comparison of species compositions also revealed a recurring pattern of asymmetry in ER–mitochondria lipid trafficking. Exclusively ER-synthesized GPLs maintained their species compositions in mitochondria (Fig. 4d,i), consistent with bulk, largely unselective transfer from the ER to mitochondria. In contrast, GPLs synthesized within mitochondria showed altered species compositions upon appearing in the ER (Fig. 4i).

The PG/BMP pool in the ER contained less PG/BMP 34:1 (32 mol% from 73 mol% in mitochondria) (Fig. 4j) and was instead enriched in PG/BMP 36:1 (26 mol%), which clustered to the ER module (Cluster V; Fig. 3d). This reshaping of species composition aligns with remodelling of mitochondria-derived PG in the ER or with selective import specific PG species into the ER.

The minor PE pool in the ER was enriched in species with two DBs (e.g., PE 34:2, 36:2) compared to the much larger mitochondrial PE pool (Fig. 4d). This distinct species composition suggests limited transfer of mitochondrial PE, keeping separation between the ER and mitochondrial PE pools produced by the two parallel pathways.

### Ethanolamine deficiency establishes the etherGPL-rich lysosomal membrane territory

Having defined general organizational principles across compartments, we next asked whether these principles are conserved across cell types or remodelled by metabolic context.

EtherGPLs differed strikingly between HCT116 colorectal carcinoma cells and their isogenic derivative HKh-2 cells (KRAS G13D knockout, KRAS^wt/–^): PC O– was strongly reduced in HKh-2 (1.4 mol% vs 8.0 mol% in HCT116), whereas PE O– remained similar (2.3 mol% vs 1.9 mol%; Fig. 5a). PAP-SL revealed a pronounced difference in lysosomes: the lysosomal levels of PC O– dropped from 17 mol% in HCT116 to 1.8 mol% in HKh-2, whereas PE O– was stable (2.4 mol% vs 2.1 mol%; Fig. 5b). This reveals that the etherGPL-enriched lysosomal membrane territory is a dynamic and tightly regulated feature of cellular lipid organization.

**Figure 5.**
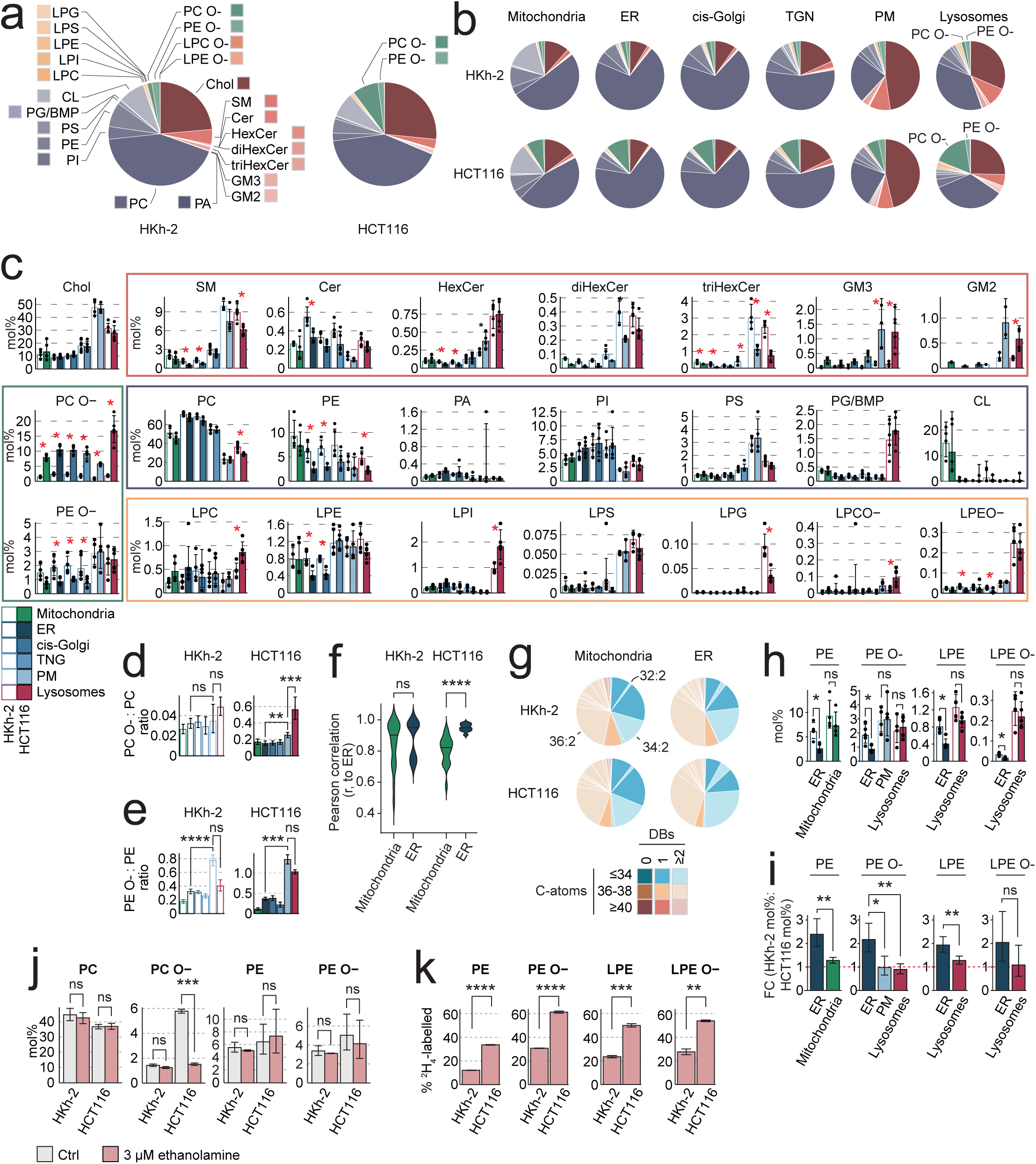
Lysosomal ether GPL territory emerges under ethanolamine deficiency. **a–b**, mol% of lipid classes in WCL (**a**) and in isolated compartments (**b**) of HKh-2 and HCT116 cells. **c**, mol% of lipid classes across compartments; asterisks denote significant differences (Welch’s t-test, FDR 10%). **d-e**, PC O–:PC (d) and PE O–:PE (e) class ratios across compartments of HKh-2 (left) and HCT116 (right). **f**, Pearson correlation coefficients (r) within ER or between ER and mitochondria based on PE species compositions. **g**, PE species compositions (mol% within class) in mitochondria (left) and ER (right) from HKh-2 (top) and HCT116 (bottom). **h**, mol% of GPL classes with an ethanolamine headgroup in the ER and other compartments from HKh-2 and HCT116 cells. i, Fold-change (FC)(HKh-2 mol% : HCT116 mol%) for indicated compartments. **j**, mol% of indicated lipid classes in WCL after 9-day supplementation with 3 µM ethanolamine. **k**, Percentage of heavy-isotope labeled lipid classes after 9-day treatment with 3 µM ^2^H_4_-ethanolamine. Statistics: Welch’s t-test with BH correction (FDR 10 %) for **c** and **h**; Welch’s t-test for **d**,**e**,**j**, and **k**. *P*<0.05, \**P*<0.01, \*\**P*<0.001, \*\*\**P*<0.0001. *N = 4–5 for a–d; N = 3 for e–f. Abbreviations as in* Fig. 1.

Both cell lines nevertheless retained the organizational principles observed in HeLa cells, including class-wise distribution patterns (Fig. 5c), the divergent trajectories of PC O– and PE O– relative to their diacyl counterparts (Fig. 5d,e), and the distinct species compositions of PE pools in the ER and mitochondria (Fig. 5f,g).

However, compartment-resolved, class-by-class comparisons revealed site-specific changes in HKh-2 accompanying the reduction in PC O-. The minor PE pool in the ER increased 2.4-fold in HKh-2 (6.1 mol% vs 2.6 mol% in HCT116), whereas the dominant pool in mitochondria showed only a modest, non-significant shift (9.2 mol% vs 7.2 mol% Fig. 5h,i). This split is consistent with enhanced PE synthesis through the CDP-ethanolamine branch of the Kennedy pathway. In line with this, other GPL classes sharing the ethanolamine headgroup with PE (PE O–, LPE, LPE O–) showed a similar ER-restricted increase: their minor pools in the ER approximately doubled, even though their major pools at the PM and in lysosomes remained unchanged (Fig. 5h,i).

Together, these patterns point to enhanced flux via CDP-ethanolamine branch of the Kennedy pathway in HKh-2 cells, offering a parsimonious explanation for the concomitant decrease in PC O– via competition for shared alkyl-DAG precursor.

Consistent with this model, supplementation with ethanolamine, which drives the CDP-ethanolamine branch of the Kennedy pathway, reduced PC O– in HCT116 cells from 5.8 to 1.6 mol% (Fig. 5j), comparable to HKh-2 cells. Importantly the supplementation did not affect PC O-levels in HKh-2 cells, suggest that these cells were already ethanolamine replete, whereas HCT116 cells operated under ethanolamine deficiency, likely reflecting their faster proliferating rate^34^. Isotope tracing further supported this: labelled ethanolamine was incorporated into 60% of PE O– in HCT116 but only 30% in HKh-2 (Fig. 5k), consistent with stronger dependence of HCT116 on exogenous ethanolamine. PE displayed the same trend, but with overall lower incorporation than PE O-, consistent with a substantial fraction of cellular PE pool being produced via PS decarboxylation.

These results demonstrate that the etherGPL-enriched lysosomal membrane territory is a dynamically regulated feature of cellular lipid organization. Its regulation is driven by PC O–, whose abundance and lysosomal enrichment depend on ethanolamine availability but remain uncoupled from diacyl PC (Fig. 5j).

Altogether, PAP-SL provides a compartment-resolved, species-level view of lipid organization in human cells. It transforms lipid atlases into organizing principles and reveals metabolic rewiring hidden in whole-cell analyses.

## Discussion

How eukaryotic cells organize thousands of lipid species to build diverse membranes and sustain organelle functions has long been a central question. Analogous to AP-MS for protein assemblies^35^, PAP-SL isolates compartments and quantifies lipid stoichiometry (mol%), generating maps of every species across the cell. Correlation analysis distills these maps into organizing rules.

First, origin sets the baseline. Lipids made in mitochondria or lysosomes remain largely confined, whereas ER-synthesized lipids disperse broadly to establish distant membrane territories. Second, specific lipid-transfer pathways carve class-wise escapes from ER to the PM: CERT delivers ceramide to the Golgi for conversion to SM^36^, and ORPs move PS and cholesterol directly to the PM^14,37^. Third, a headgroup-agnostic, species-level selection further sculpts these flows: more saturated GPL species across multiple headgroups are preferentially retained at the PM, likely reflecting favorable interactions with SM and cholesterol^6^. Together, these layers explain how a cholesterol/SM/PS-rich and mechanically robust PM is maintained. ER supplies mitochondria with less saturated GPLs in bulk and non-selectively, consistent with lipid-transfer bridges/tethers such as VPS13A and the ER–mitochondria tether PDZD8^38–40^. Finally, ER establishes a distinct lysosomal territory enriched in ether-linked GPLs.

PAP-SL revealed yet another layer, emerging from the ether linkage. Despite sharing the Kennedy pathway with their diacyl counterparts, ether linkage directed GPLs toward a lysosomal membrane territory. This pattern could reflect direct ER– lysosome transfer at contact sites where LTPs such as VPS13C operate^38,39^, but a more likely explanation is selective persistence: etherGPLs are generally poorer substrates for lysosomal phospholipases and the ether bond is chemically harder to hydrolyze^33,41^. Persistence of PC O-may stabilize lysosomal membranes against hydrolytic stress and reduce futile degradation–resynthesis cycles in fast-growing cells such as HCT116.

PE O-followed a different route. It was abundant at both PM and lysosomes but showed no further gain across the PM-to-lysosome transition. Ether linkage alone is thus insufficient to predict lysosome-specific buildup. A plausible biochemical asymmetry is that TMEM189 preferentially introduces a vinyl-ether bond into PE O-^41^, consistent with its high unsaturation; this bond can be targeted by specific turnover enzymes such as TMEM86A and cytochrome c^42,43^, and may also promote tighter membrane packing^44^, offering a physical rationale for its PM enrichment. Consistent with this idea, a recent study using bifunctional ether-lipid probes reported slower intracellular movement for vinyl-ether versus alkyl-ether species^45^. Topology may also contribute. Like PE, PE O-is comparatively enriched on the cytosolic leaflet of the PM^9^, leaving it cytosolic during endocytosis, shielded from luminal hydrolases, and available for retrieval through contact-site transport—limiting further lysosomal accumulation.

Finally, compartment-resolved comparisons expose regulation hidden in whole-cell lipidomics. The lysosomal PC O-territory behaved as a marker of metabolic state, collapsing when ethanolamine supply/flux shifted, while PC (diacyl) remained stable— consistent with specific competition for alkyl-DAG in the CDP-ethanolamine branch and with ethanolamine demand scaling with fast growth of HCT116^34^.

Together, PAP-SL delivers a species-level atlas of lipid distributions from matched preparations and moves beyond mapping to extract rules of organization. This steady-state framework complements perturb-and-observe approaches such as photo-caged lipid probes or quantitative imaging of lipid transport, providing the spatial context for interpreting acute manipulations^46,47^. PAP-SL therefore opens the way to address how lipid organization is dynamically rewired across cell states and in disease.

## Acknowledgements

The research reported in this publication was supported by Danish National Research Foundation (DNRF125 to M.J.), The Independent Research Fund Denmark (6108-00542B to K.M), Novo Nordisk Foundation (NNF15OC0018914 and NNF24OC0096144 to M.J.), and Danish Cancer Society (R124-A7929-15-S2 to K.M). We are thankful to the Lipidomics Core Facility at the Danish Cancer Institute for access to instrumentations.

## Methods

### Cell lines

HeLa cells were purchased from ATCC (Manassas, VA, USA). HCT116 and its isogenic derivative HKh-2 (KRAS G13D allele knockout, KRAS^wt/–^)^34^ was a kind gift from Dr. S. Shirasawa. Dulbecco’s Modified Eagle Medium (DMEM; cat. no 31966021), TrypLE™ Express (cat. no 12604013), and Dulbecco’s Phosphate Buffered Saline (PBS; cat. no 14190094) were purchased from Gibco (Carlsbad, CA, USA). Anti-Rabbit IgG MicroBeads (cat. no 130-048-602), Streptavidin MicroBeads (cat. no 130-048-102), and MS columns (cat. no 130-042-201) were purchased from Miltenyi Biotec (Bergisch Gladbach, Germany).

### Parallel (immuno)affinity purification of cellular compartments

Cells were cultured to 90% confluence in 15-cm dishes (one dish per compartment) in Dulbecco’s Modified Eagle Medium (DMEM; Gibco, Carlsbad, CA) supplemented with 6%(v/v) heat-inactivated fetal calf serum (FCS; Gibco). Cells were maintained at 37 °C in a humidified incubator with 5% CO₂. Cells in an additional dish were used for cell counting with a hemocytometer.

Cell surface biotinylation was performed on two dishes of cells prior to harvesting. Cells were washed three times in ice-cold phosphate-buffered saline (PBS; Gibco) and incubated for 30 min at 4 °C with 5 ml PBS containing 1 mg/ml EZ-Link Sulfo-NHS-LC-LC-Biotin (Thermo Fisher Scientific, cat. no. 21338). After three washes in PBS, biotinylated cells were pooled with untreated cells for downstream processing. Cells were harvested by scraping in PBS, washed in SuMa buffer [10 mM HEPES, 0.21 M mannitol, 70 mM sucrose, pH 7.5], and resuspended in SuMa4 buffer [SuMa buffer + 0.5 mM DTT, 0.5 % BSA (essentially FA-free, A3803, Sigma Aldrich), 25 units/ml Benzonase (E1014; Sigma Aldrich) and 1x cOmplete™ Mini, EDTA-free Protease Inhibitor Cocktail (04693159001, Roche Diagnostics)] to 60 x 10^6^ cells per ml. HeLa cells were lysed in a gentleMACS™ Dissociator (130-093-235; Miltenyi Biotech). HCT116 and HKh-2 cells were lysed by syringe passage (25G needle, 3 or 7 passes, respectively). Post-nuclear fractions (PNFs) were prepared from whole-cell lysate (WCLs) by three cycles of centrifugation at 1000 g, for 5, 10, and 15 min, 4 °C, with the supernatant transferred to new tubes after each cycle.

Cellular compartments were purified using antibody- and streptavidin(PM)-based capture. PNF aliquots (equivalent to 9×10⁶ cells) were incubated with primary antibodies (0.18 µg) for 1 h at 4 °C: rabbit anti-Tom20 (Abcam, ab78547) for mitochondria, rabbit anti-Calnexin (Abcam, ab22595) for ER, rabbit anti-RCAS1 (Cell signaling technology, 12290S) for cis-Golgi, rabbit anti-TGN46 (Abcam, ab16052) for TGN, rabbit anti-LAMP1 (Abcam, Ab24170) for lysosomes, rabbit anti-Pex14 (Proteintech, 10594-1-AP) for peroxisomes, or a negative control antibody (Rabbit IgG Isotype Control; Thermo Fisher Scientific, 31235)(no antibody added for PNF for PM purification). PNFs were further incubated for 1 h with 25 µl Anti-Rabbit IgG MicroBeads (cat. No 130-048-602; Miltenyi Biotec, Bergisch Gladbach, Germany), or, for PM capture, 25 µl Streptavidin MicroBeads (cat. No 130-048-102; Miltenyi Biotec). PNFs were then loaded on pre-equilibrated (with 500 µl SuMa4 buffer) MS columns on a magnetic stand, washed once with 500 µl SuMa4 buffer, once with 500 µl SuMa2 buffer (SuMa4 without Benzonase and protease inhibitor), and once by filling up the reservoir (∼3.5 ml) with SuMa2 buffer. The bound materials were eluted in 600 µl SuMa2 buffer using the supplied plunger. Eluates were divided into fractions of 150 µl for western blotting and fractions of 450 µl for lipidomics. All samples were pelleted at 21,100 g for 20 min at 4 °C. Lipidomics samples were stored at -80 °C and western blot samples were dissolved in 1x SDS-PAGE sample buffer [62.5 mM Tris, 70 mM sodium dodecyl sulfate (SDS), 0.15 mM bromophenol blue, 10% (v/v) glycerol, pH 6.7] containing 100 mM DTT and stored at -20 °C. WCLs and PNFs equivalent of 450,000 cells were added to 450 µl (for lipidomics) or 150 µl (for Western blotting) SuMa4 buffer and prepared parallel with the eluates.

### Lipidomics

Lipids were extracted and analyzed as described previously^26^. Briefly, pellets were mixed with internal lipid standards (see Extended Data Table 1) and chloroform:methanol (2:1, v/v), vortexed, centrifuged, and washed sequentially. Lower organic phases were collected, dried in a vacuum centrifuge, and resuspended in chloroform:methanol (1:2, v/v). Shotgun lipidomics was performed using a Q Exactive Hybrid Quadrupole–Orbitrap mass spectrometer (Thermo Fisher Scientific) equipped with a TriVersa NanoMate (Advion Biosciences, Ithaca, NY). Both positive- and negative-ion modes were recorded in technical duplicates. Blank samples containing only internal standards were processed in parallel.

For supplementation with 3 µM ethanolamine (cat. No 411000; Sigma-aldrich) or 3 µM 1,1,2,2-D₄-ethanolamine (catalog no. DLM-552-0.1, 98 %, Eurisotop/Cambridge Isotope Laboratories, Paris, France), cells were cultured as above, but in 24-well plates, for 9 days with or without the supplement, and analyzed with shotgun lipidomics as described above. Species compositions of PG and BMP were determined on HeLa cells cultured in 24-well plates and analyzed using LC–MS as previously^26^.

### Lipid nomenclature

Lipids are annotated according to the LIPID MAPS classification system^1,2,48^ with the alteration that the lipid category GPL has been divided into acyl-acyl GPLs (diacylGPLs), alkyl-acyl GPLs (etherGPLs), and monoacyl and monoalkyl GPLs GPLs (lysoGPLs). Lipid species are reported according to their sum composition, e.g., PC 34:1, belonging to the PC class and having 34 total acyl-C-atoms and 1 total acyl DB. For SLs, number of OH groups are indicated after semicolon (only sphingolipids with 2 OH groups were quantified in this study).

### Lipidomics data processing

Extraction of ion intensities and *m/z* values and lipid identification were carried out in LipidXplorer version 1.2.4^49,50^ using class-specific fragmentation criteria (Extended Data Table 2) defined in MFQL (Molecular Fragmentation Query Language) files. Quantification was performed by internal standard normalization with LipidQ (https://github.com/ELELAB/lipidQ and aggregated by the means of technical duplicates. Species not passing blank filtering (abundance <5 × blank or <0.001 pmol), replicate filtering (detection in at least 4 out of 5 experiments), or database filtering (C-atoms and DBs: 14:0, 16:0-1, 18:0-3, 20:0-5 and 22:0-6 (in individual chains) in GPLs and 32-44:0-3 (in total) in SLs) were excluded (see Extended Data Fig. 2 for filtered species).

### Data analysis and statistics

Data were expressed as molar amounts (pmol), molar percentages of the total lipidome (mol%), or molar percentages within classes (mol% of class). Principal component analysis (PCA) was performed using log_2_-transformed mol% of species and the prcomp() function in R version 4.2.3^51^, with missing values set to 0.00001 mol%. Pearson correlation coefficients (r) between species were calculated on mol% using cor() function of R, setting missing values to 0, and correlation heatmaps were (unsupervised) hierarchically clustered and visualized using the pheatmap() function from the R package pheatmap version 1.0.12^52^. For network visualization, lipid species with Pearson’s *r* > 0.7 or < –0.7 were exported to Cytoscape v3.9.1 and displayed using an edge-weighted spring-embedded layout. Species without such correlations, as well as isolated small networks (≤2 species), were excluded.

For correlation analysis of species compositions across compartments, Pearson *r* values were calculated from log2(mol% of class) and visualized as heatmaps using *Morpheus* (https://software.broadinstitute.org/morpheus/). For comparison of ER–ER vs ER–other compartments correlations, *r* values were Fisher Z-transformed and compared statistically with Welch’s ANOVA with Benjamini and Hochberg (BH) correction^53^. Cosine similarities of species compositions (mol% of class) of classes were calculated in Microsoft Excel (for Microsoft 365, v16.0; Microsoft, Redmond, WA, USA) and hierarchically clustered (1-Cosine similarity) and visualized in *Morpheus*. For comparison of mol% of lipid classes within each compartment between HKh-2 and HCT116, values were log₂-transformed and tested with Welch’s t-test with BH^53^ correction. When a lipid class was detected in only one cell line for a given compartment, significance was assessed by a one-sided one-sample t-test with the null hypothesis mean ≤ log₂(0.1). Figures were generated using the R package *ggplot2* v3.4.0^54^, GraphPad Prism v10.6.0, and Adobe Illustrator. Mean and standard deviation in bar graphs were calculated from geometric values.

### Western blotting

Proteins were separated on 4–15% PROTEAN® TGX™ Precast Protein Gels (cat. No 4561093; Bio-Rad) and transferred to nitrocellulose membrane using a Trans-Blot Turbo Transfer System (Bio-Rad). Membranes were incubated with primary antibodies against: EEA1 (rabbit, cat. No ab2900; Abcam), histone H3 (rabbit, cat. No ab18521; Abcam), Tom20 (rabbit, cat. No sc11415; Santa Cruz), cytochrome c (mouse, cat. No 33-8500; Invitrogen), VDAC (rabbit, cat. No 12454; Cell Signaling Technology), Calnexin (mouse, cat. No sc-80645; Santa Cruz), PDI (rabbit, cat. No Ab3672; Abcam), RCAS1 (rabbit, cat. No 12290S; Cell Signaling Technology), TGN46 (rabbit, cat. No ab16052; Abcam), LAMP2 (mouse, cat. No H4B4; Developmental Studies Hybridoma Bank), Cathepsin B (mouse, A kind gift from Dr. Ekkehard Weber (Martin Luther University Halle-Wittenberg, Halle, Germany)), Pex14 (rabbit, cat. No 10594-1-AP; Thermo Fisher Scientific), and Catalase (rabbit, cat. No ab209211; Abcam).

Secondary antibodies included horseradish peroxidase (HRP)-conjugated rabbit anti-mouse antibody (cat no. P0260; Dako), mouse anti-rabbit light-chain antibody (cat no. 211-032-171, Jackson ImmunoResearch) and native protein A (ab7456; Abcam). Biotinylated proteins were detected with Streptavidin-HRP (ab7403; Abcam). Membranes were developed with Clarity Western ECL Substrate (170-5061; Bio-Rad) using a LAS-3000 imaging system (Fujifilm, Tokyo, Japan).

---

**Extended Data Figure 1.**
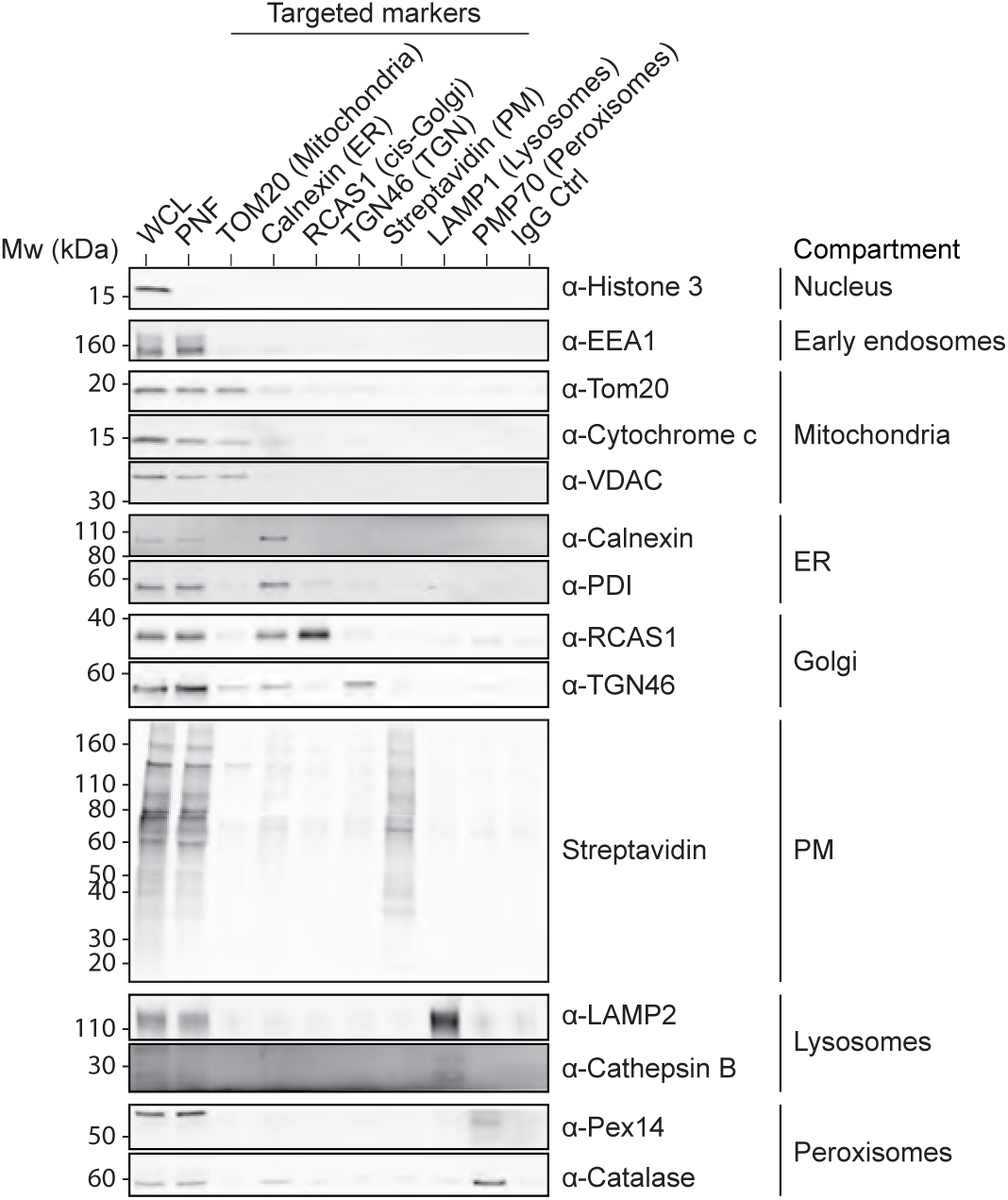
Western blot evaluation of PAP-SL fractions from HeLa cells. The figure shows a representative image.

**Extended Data Figure 2.**
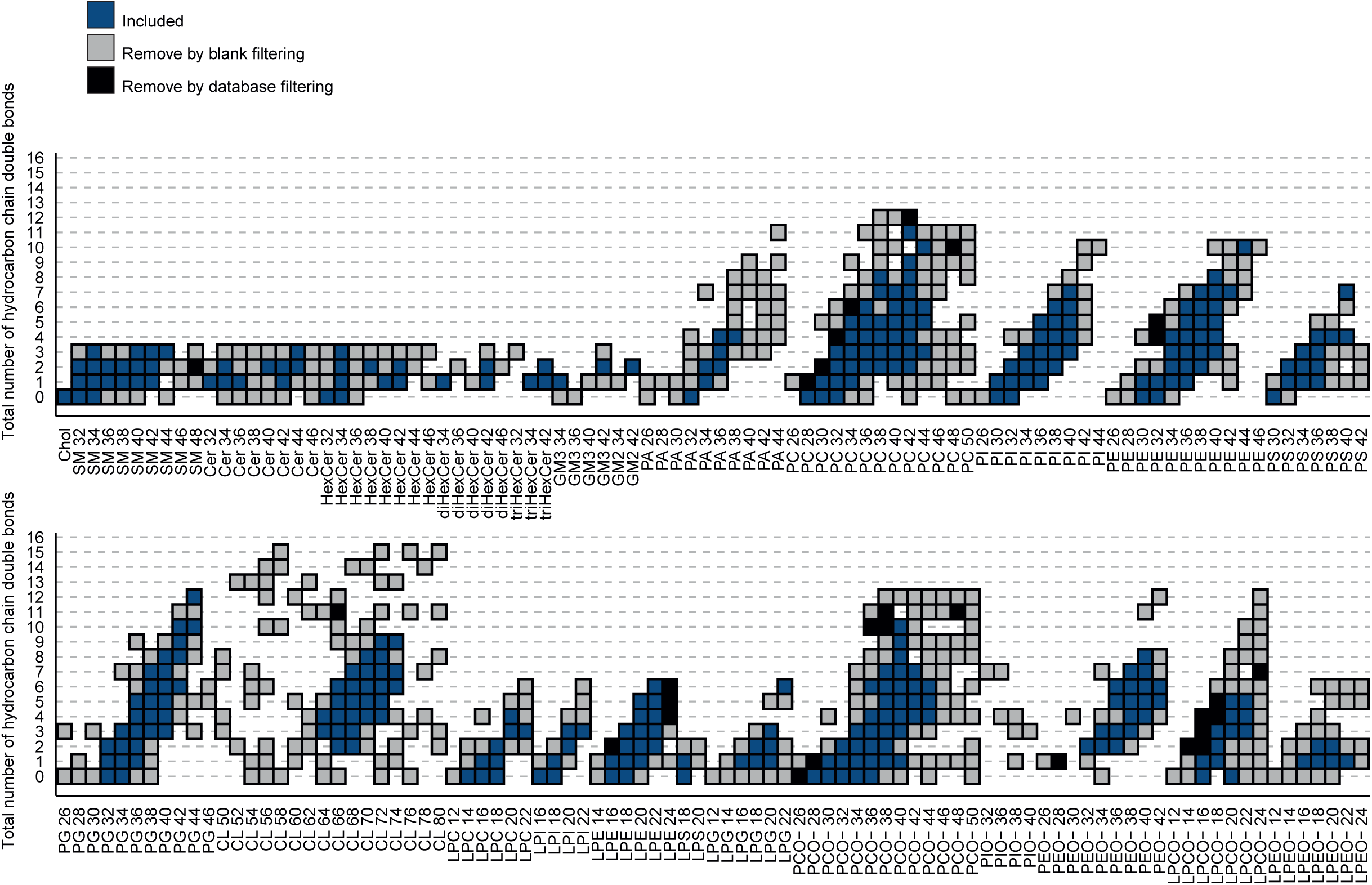
Filtration of detected lipid species. Lipid species are indicated with their classes and total numbers of hydrocarbon chain carbon atoms (horizontal) and double bonds (vertical). Detected lipid species were either included in the final datasets or filtered via blank filtering or database filtering (see Methods).

**Extended Data Figure 3.**
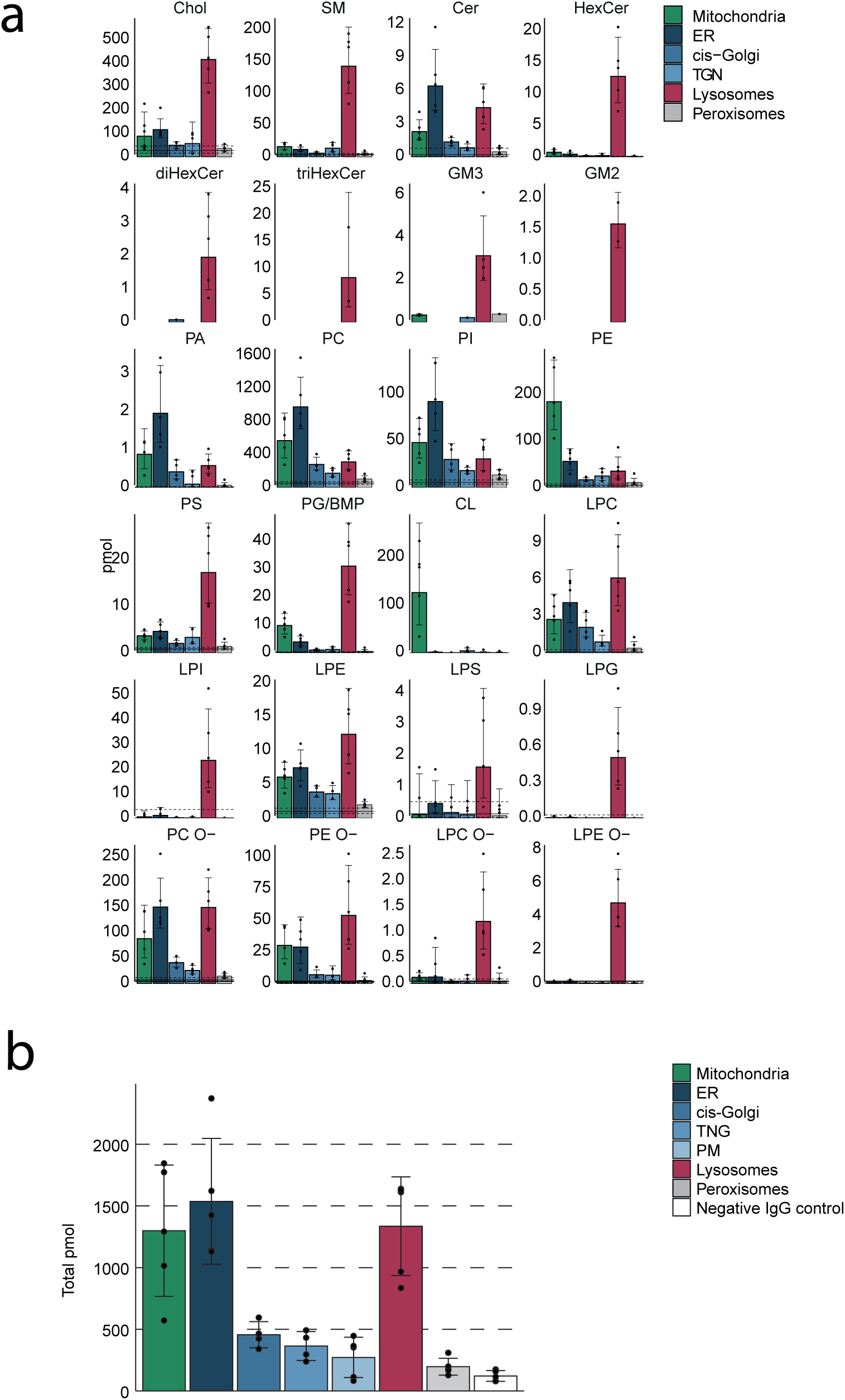
Absolute quantities of lipids in PAP-SL fractions from HeLa cells. **a**, Absolute quantities of lipid classes detected in cellular compartments from HeLa cells. Bar graphs show mean and standard deviations from 4-5 independently performed experiments. The horizontal line indicates mean for negative IgG control, and the dashed lines standard deviations. **b**, Absolute total quantities of lipids in isolated fractions. Bar graphs show mean and standard deviations from 4-5 independently performed experiments.

**Extended Data Figure 4.**
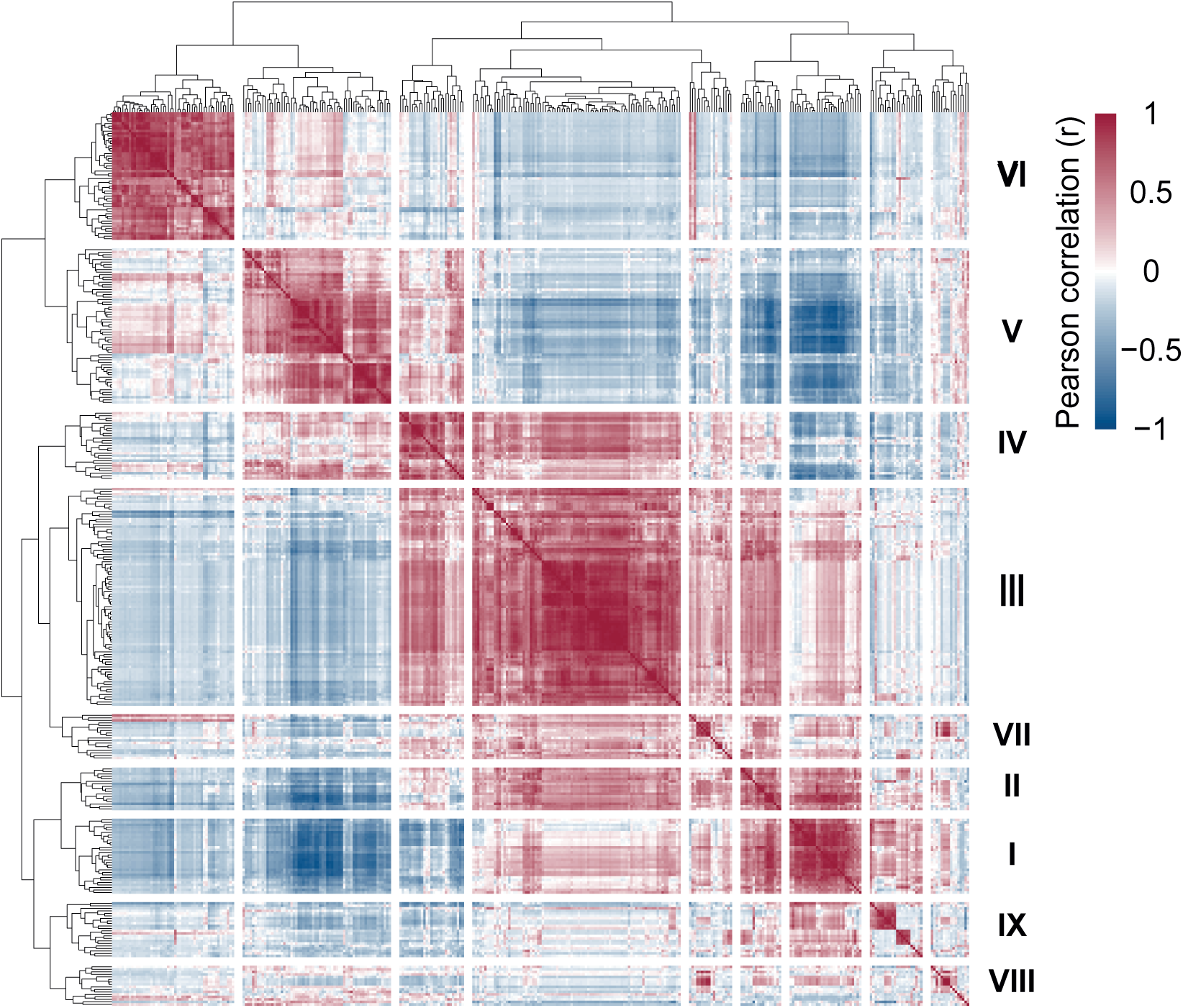
Correlation matrix of lipid species in HeLa cells. Pearson correlation coefficients were calculated for all pairs of lipid species and the matrix was hierarchically clustered.

**Extended Data Table 1.**
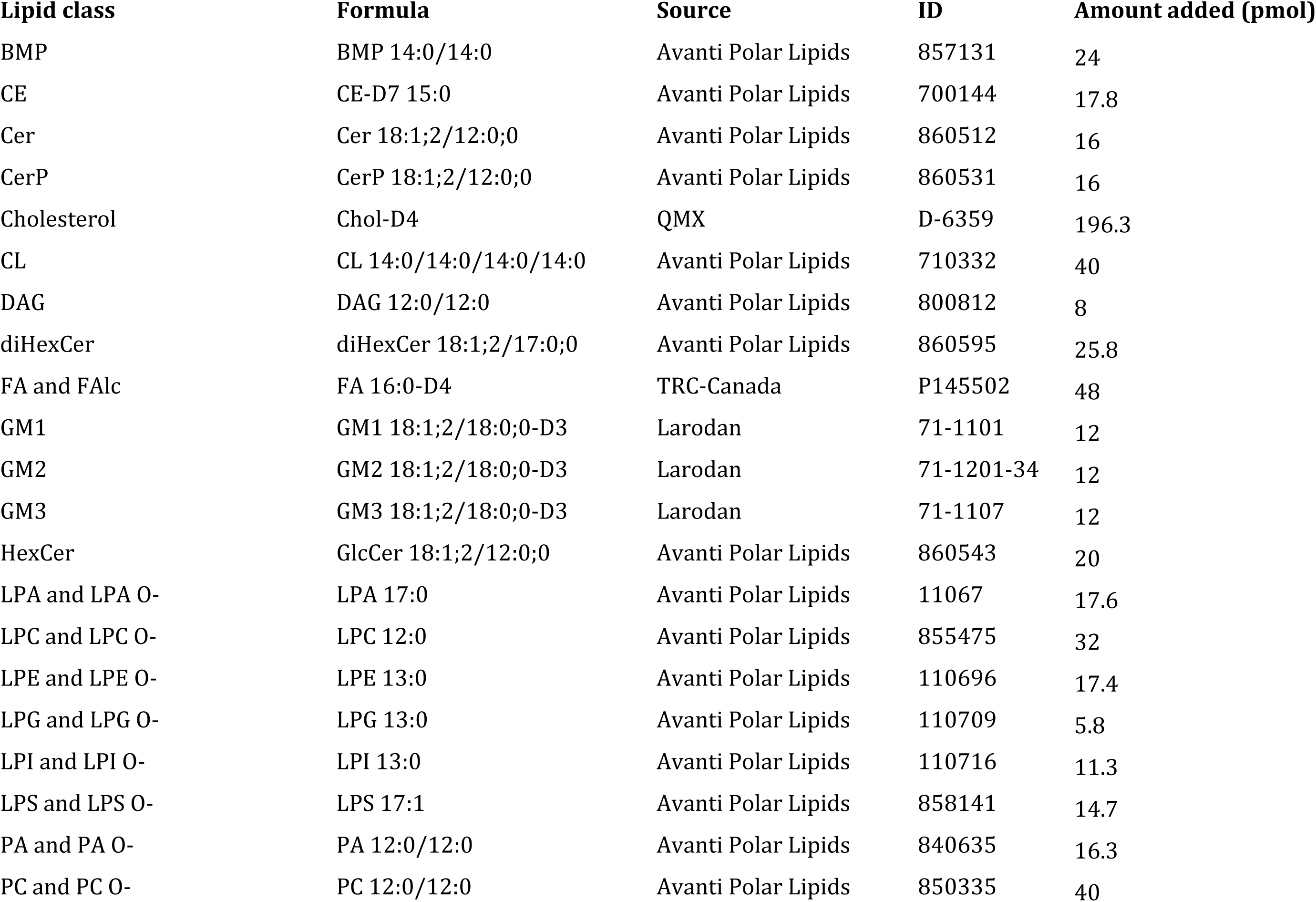
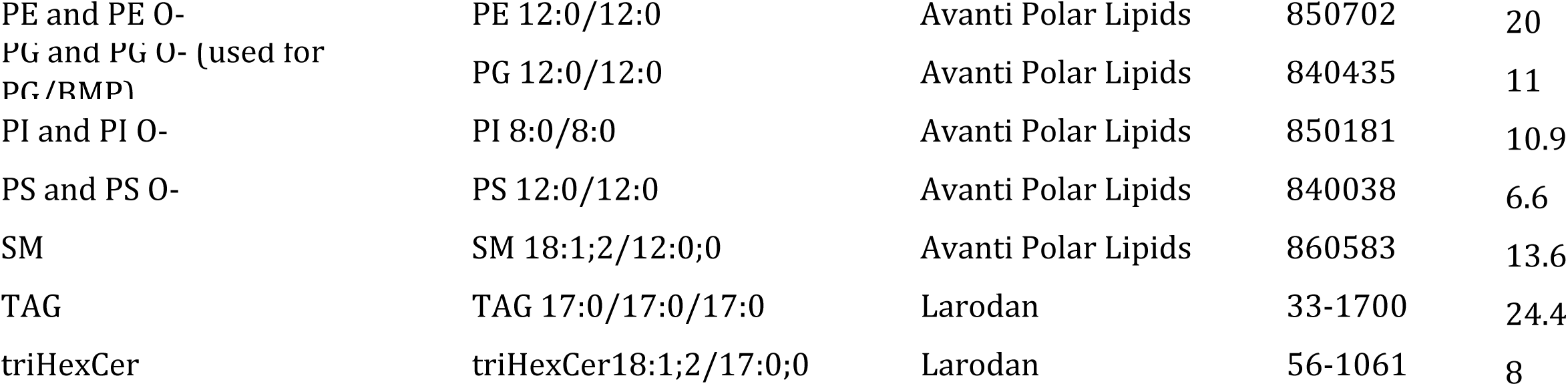
List of internal lipid standards.

**Extended Data Table 2.**
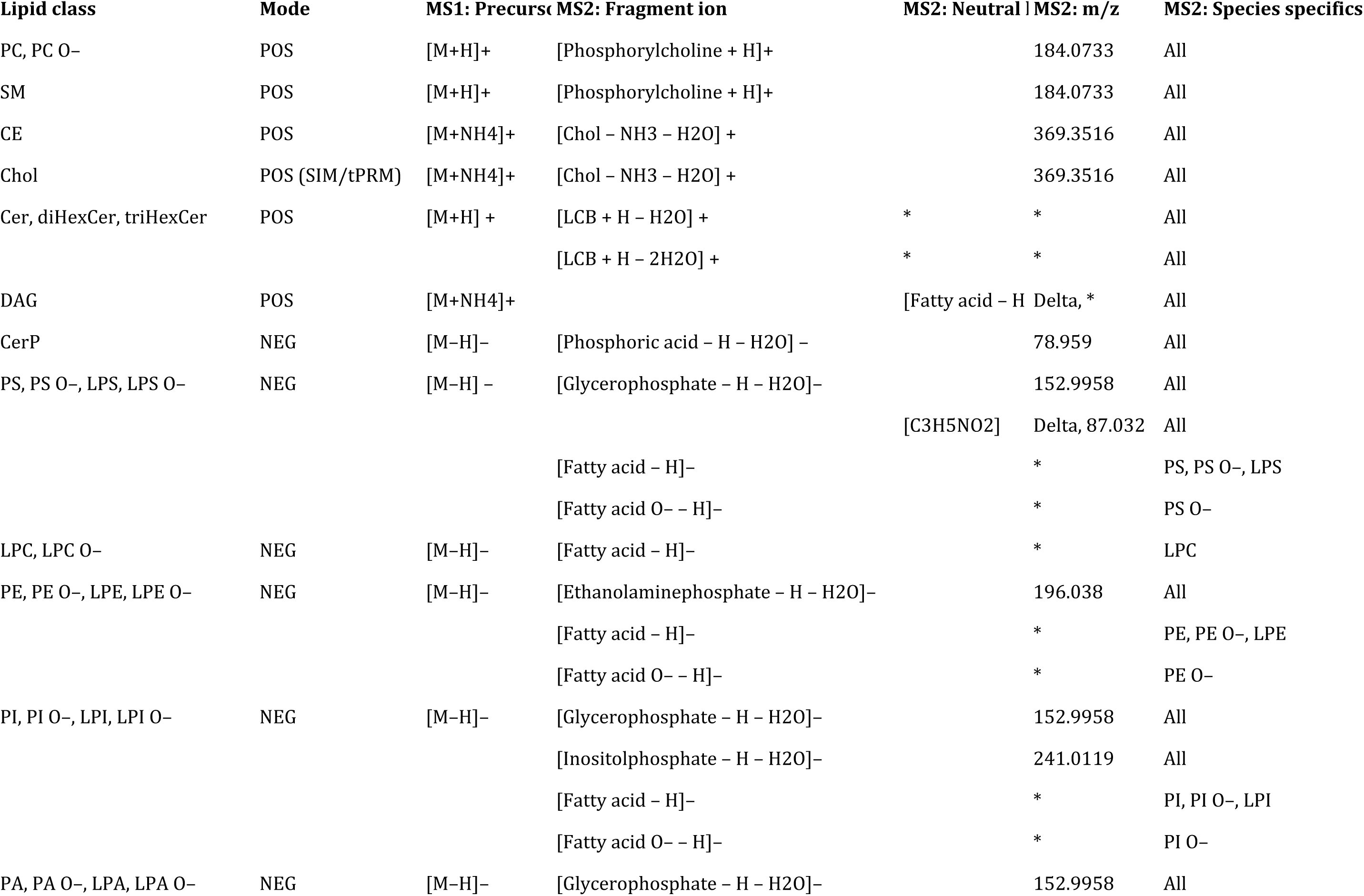
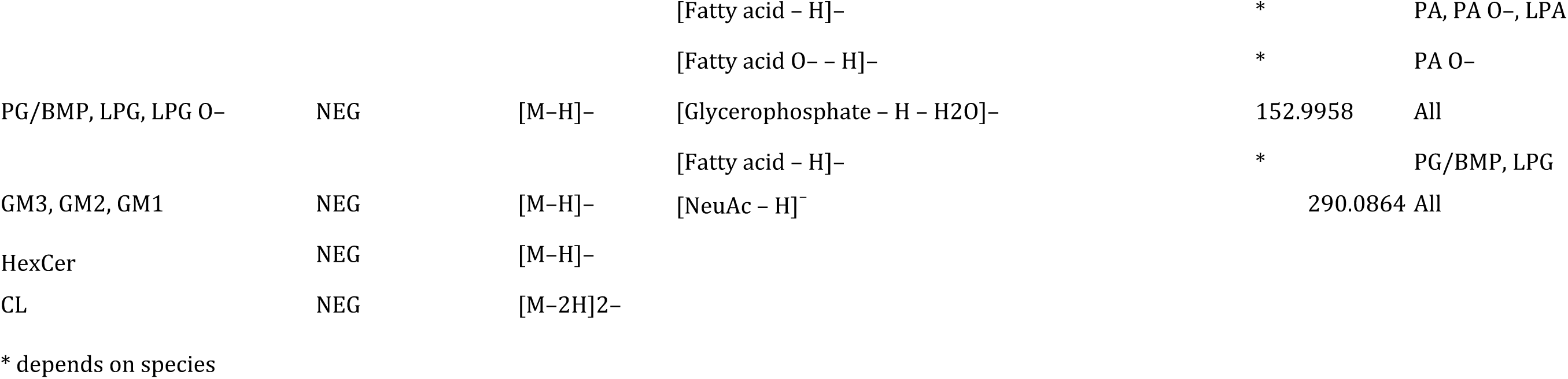
List of ions and neutral losses uses as identification criteria for lipid classes.

